# Tuning and validation of a virtual mechanical testing pipeline for condylar stress fracture risk assessment in Thoroughbred racehorses

**DOI:** 10.1101/2024.10.27.620476

**Authors:** Soroush Irandoust, R. Christopher Whitton, Corinne R. Henak, Peter Muir

**Affiliations:** Department of Surgical Sciences, School of Veterinary Medicine, University of Wisconsin-Madison, Madison, WI 53706, USA; Department of Mechanical Engineering, University of Wisconsin-Madison, Madison, WI 53706, USA; Equine Centre, Melbourne Veterinary School, University of Melbourne, Werribee, Vic, 3030, Australia; Department of Biomedical Engineering, University of Wisconsin-Madison, Madison, WI 53706, USA; Department of Orthopedics and Rehabilitation, University of Wisconsin-Madison, Madison, WI 53705, USA

**Keywords:** Thoroughbred racehorses, third metacarpal bone, condylar stress fracture, functional adaptation, finite element analysis

## Abstract

Condylar stress fracture of the third metacarpal/metatarsal bone (MC3/MT3) in Thoroughbred racehorses is a common catastrophic injury, putting racehorse and jockey safety at risk. Microdamage forms in the distal MC3 and may result in strain elevation and higher risk of stress fracture, directly and through focal osteolysis resulting from the associated process of repair by remodeling in the condylar parasagittal grooves (PSG). Standing computed tomography (sCT) is a practical screening tool for detection of fatigue-induced structural changes, but the clinical interpretation remains subjective. The goal of this study was to develop and validate a sCT-based subject-specific finite element analysis (FEA) pipeline for virtual mechanical testing of the distal MC3. Twelve (n=12) MC3 condyles with available *ex vivo* strain were selected. Half of the specimens were used for tuning of the Young’s modulus in fatigue sites, and the other half to examine validity of the pipeline for prediction of PSG strain. The tuned model improved prediction of strain and predicted higher strain in bones with PSG lysis over the untuned model, which is an essential feature for a diagnostic screening tool. The presented pipeline is expected to assist clinicians with interpretation of sCT-detectable structural changes and ultimately condylar fracture risk assessment.

## 1. Introduction

Condylar stress fracture of the metacarpophalangeal/metatarsophalangeal (MC3/MT3, fetlock) joint is a common catastrophic injury in Thoroughbred racehorses worldwide and a major cause of euthanasia [1–3]. It is the second most common serious musculoskeletal injury in the Unites States after proximal sesamoid bone (PSB) fracture [4,5]. In addition to racehorse injury, jockey falls and injuries are most associated with fetlock injuries many of which are serious and lead to life-changing disability. Jockeys were ∼160-200 times more likely to fall and get injured when they rode a horse that went on to sustain catastrophic musculoskeletal injury compared with riding an uninjured horse [6,7]. Focal microdamage, incomplete fractures, and preexisting resorption at sites prone to condylar fracture have been observed in the fractured and contralateral nonfractured limbs of Thoroughbred racehorses [8–15] confirming that condylar fractures of the MC3/MT3 are fatigue-induced stress fractures [11]. Thoroughbreds with a complete displaced condylar stress fracture have a significantly decreased prognosis for racing after surgery [16–19], hence, prevention of these injuries and associated horse death and jockey injury from falls is an important focus.

Large cyclic loads and microdamage accumulation in the distal end of the MC3/MT3 trigger new trabecular bone modeling and subchondral sclerosis [20,21], which is thought to help unload fatigued bone, allowing focal remodeling to repair damage. Site-specific microdamage in bone can be resorbed through bone remodeling, but because repetitive high loads suppress bone resorption, damage accumulation often outpaces this process [22]. Such an imbalance can result in subchondral bone fatigue injury (SBI) in the parasagittal grooves (PSG), visible as focal osteolysis lesions with diagnostic imaging. Both focal microdamage and osteolysis have been shown to compromise distal metacarpal subchondral bone mechanical properties *ex vivo* [23,24].

sCT is a practical noninvasive racehorse screening method with high sensitivity for detecting structural changes associated with fatigue damage [25–27]. Despite high sensitivity of sCT for detection of these changes, clinical interpretation is still challenging and an objective classifier for condylar fracture risk assessment is lacking. Thus, combining sCT with physics-based predictive models has the potential to improve identification of MC3/MT3 bones at risk of fracture.

CT-based subject-specific finite element (FE) modeling is a helpful tool for virtual noninvasive mechanical testing and has been extensively used for biomechanical assessment of human hips and spines [28–30]. In the fetlock of Thoroughbred racehorses, although prior microCT-based FE analysis (FEA) has shed light on the biomechanical properties of the subchondral bone [31], there is currently an urgent need for a sCT-based FEA pipeline to assist clinicians with condylar fracture risk assessment. Therefore, the goal of this study was to develop and validate a sCT-based subject-specific FEA pipeline for virtual mechanical testing of the distal MC3 in Thoroughbred racehorses.

## 2. Materials and Methods

### 2.1. Specimen selection

A library of twelve MC3 condyles (n=12) from eleven thoracic limb specimens, with available surface strain data under quasi-static loading of these condyles from previous *ex vivo* mechanical testing [23], were used for this study. Specimens were collected from Thoroughbred racehorses that were euthanatized because of catastrophic racetrack injury. Age, sex, and training history of the specimens were not available. These condyles included specimens with (n=4, PSG-SBI group) and without (n=8, CTRL group) sCT-detectable focal lysis in the PSG of the distal end of the MC3 bone. Existence of fatigue damage in the PSG of the specimens in the PSG-SBI group was previously confirmed following soft tissue dissection [23,25].

### 2.2. sCT acquisition and calibration

The sCT scans of the frozen limbs were acquired using an Asto CT (Middleton, WI, USA) Equina^®^ scanner at exposure of 160 kVp and 8 mA, with 0.55 mm slice thickness. An electron density phantom (model 062M, CIRS Inc, Arlington, VA) with 4 calcium hydroxyapatite (HA) plugs with densities of 200, 800, 1250, and 1750 mg/cm^3^ was scanned with the same scanner at the same exposure settings to find the equivalent radiological or CT density, *ρ*_*CT*_(*mg_HA_/cm*^*3*^, **Figure S1**):

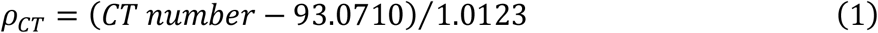

### 2.3. Segmentation of the distal MC3

Mimics (v.26) was used to segment the distal MC3 (**Figure 1**). Using a gradient-based segmentation tool, the external surface of the bone was detected semi-automatically for all individual sCT image slices sequentially. After creating the 3D distal MC3, using morphology operations one pixel was eroded from the external surface to make sure it was tight on the cortex before smoothing and importing it into 3-matic (v.18) for further processing. An anatomical coordinate system was established by first finding the long axis of the bone using a cylinder fit. The transverse plane was then constructed perpendicular to the long axis of the bone. The sagittal plane was established perpendicular to the transverse plane through the dorsal and palmar aspects of the sagittal ridge. The frontal plane perpendicular to the transverse and frontal planes was then constructed. The distal 2.5 inches of the MC3 was isolated and a homogenous triangular mesh with 0.25 mm edge length was generated.

**Figure 1.**
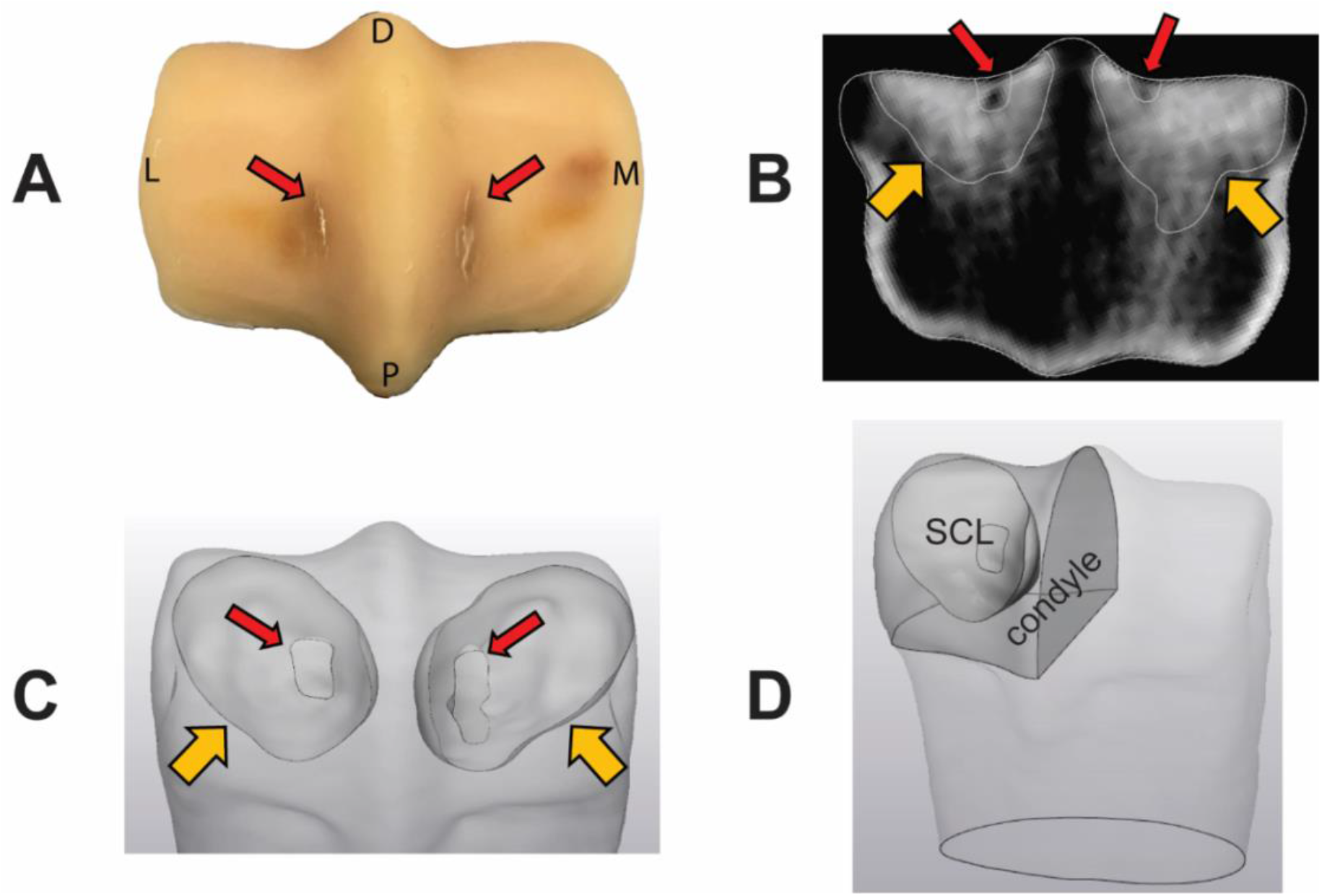
Segmentation of the sclerotic and lytic regions. Adaptive and damage factors were used for tuning of Young’s modulus in sclerotic and lytic regions indicated with orange (wide) and red (thin) arrows, respectively. Fatigue cracks are visible on the palmar aspect of the parasagittal grooves of the condyle PSG-SBI-1 after dissection and removal of soft tissue **(A)**. Sclerotic and lytic regions of PSG-SBI-1 were segmented from the standing CT image **(B, C)**. A representation of measurement of the volume of a condyle with its sclerotic region is shown **(D)**.

### 2.4. Density-Modulus relationship

Mechanical properties of bone have a strong correlation with its density [32], and this relationship can be used to predict heterogenous bone mechanical properties. First, CT density from equation (1) was converted to ash density using the data from Knowles et al. [33]:

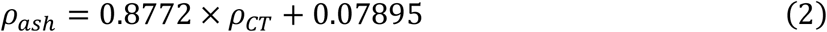

In equation (2) *ρ*_*ash*_ and *ρ*_*CT*_ are expressed in *g/cm*^*3*^. The Young’s modulus was then calculated using the experimental data collected from horse limbs by Moshage et al. [34]:

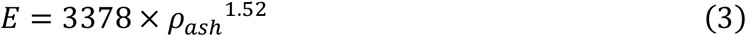

In equation (3) *E* is expressed in *MPa* and *ρ*_*ash*_ in *g/cm*^*3*^.

### 2.5. Modulus tuning for the subchondral sclerotic and lytic regions

Using the available *ex vivo* mechanical testing data, maximum principal strain in the PSG at the peak displacement was compared against FEA-predicted strain in the PSG. In our preliminary investigations we found that FE predictions of PSG strain using the density-modulus relationship in equation (3) does not match well with the experimental results. This was likely due to the complex mechanical properties of fetlock subchondral bone in actively training and racing Thoroughbreds. Microdamage is not captured with clinical CT due to resolution limitations, and although density-modulus relationship for the subchondral equine MC3 bone have been reported before at the micro scale [31], there is no available data at the macro scale. To address these two main limitations, sclerotic and lytic regions, if present, were segmented separately (**Figure 1**). To separate these regions from the rest of the distal MC3, a threshold of *HU* = 1500, equivalent to 1390 *mg_HA_/cm*^*3*^, was used based on the location of the secondary peaks in the voxel density histograms of the distal MC3 that indicate the adaptive response, as previously discussed [23]. Trabecular regions with *HU* > 1500 were defined as the sclerotic region (SCL), and isolated focal regions in the PSG with *HU* < 1500, if present, were defined as the lytic region (LYS). A modified density-modulus relationship was used by incorporating an adaptive factor for the sclerotic region (−1 < *A*_*SCL*_ < 0.5) and a damage factor for the lytic region (0.5 < *D*_*LYS*_) as shown in equations (4) and (5).

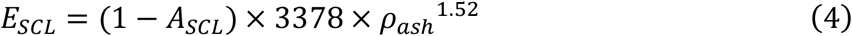

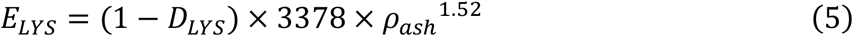

To find the optimal *A*_*SCL*_, the normalized volume of the sclerotic region was measured 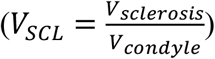 and half of the CTRL condyles were chosen (n=4) in such a way to include condyles with small and large sclerotic volumes and low and high *ex vivo* PSG max principal strain. For each of the four models, all *A*_*SCL*_ values between -1 and 0.5 were used at 0.05 intervals to find the optimal value that resulted in an average PSG max principal strain closest to the experimentally measured values. Since it has been shown that bones with larger sclerotic volumes contain higher degrees of microdamage in the subchondral bone [21], the optimal *A_SCL_* was tuned as a function of *V*_*SCL*_ (**Figure 1**).

To find the optimal *D*_*LYS*_, half of the condyles with PSG SBI (n=2) were selected and all *D*_*LYS*_ values between 0.5 and 0.95 were used at 0.05 intervals to find the optimal value that resulted in an average PSG max principal strain closest to the experimentally measured values. Due to the limited number of condyles in this group (n=2), the average of the optimal *D*_*LYS*_ for these two limbs was determined as the optimal value. The calibrated *A_SCL_* was used for defining the Young’s modulus in the sclerotic regions in these limbs.

### 2.6. Discretization, loading and boundary conditions

FE models were created in FEBio Studio (v.2.3) with 0.25 mm edge length linear tetrahedral elements in the osteolytic and sclerotic regions and 1 mm edge length linear tetrahedral elements elsewhere. Adequate mesh density was confirmed via mesh convergence analysis (**Figure S2**). A Poisson’s ratio of 0.3 and sCT-based Young’s modulus were assigned to individual elements using the density-modulus relationships detailed in sections 2.4 and 2.5.

The proximal end of the model was fully constrained and a 7.5 kN load was applied to the palmar surface of the medial condyle at 60 and 30 degrees with respect to the frontal and transverse planes, respectively, replicating the experimental loading conditions [23] (**Figure 2**).

**Figure 2.**
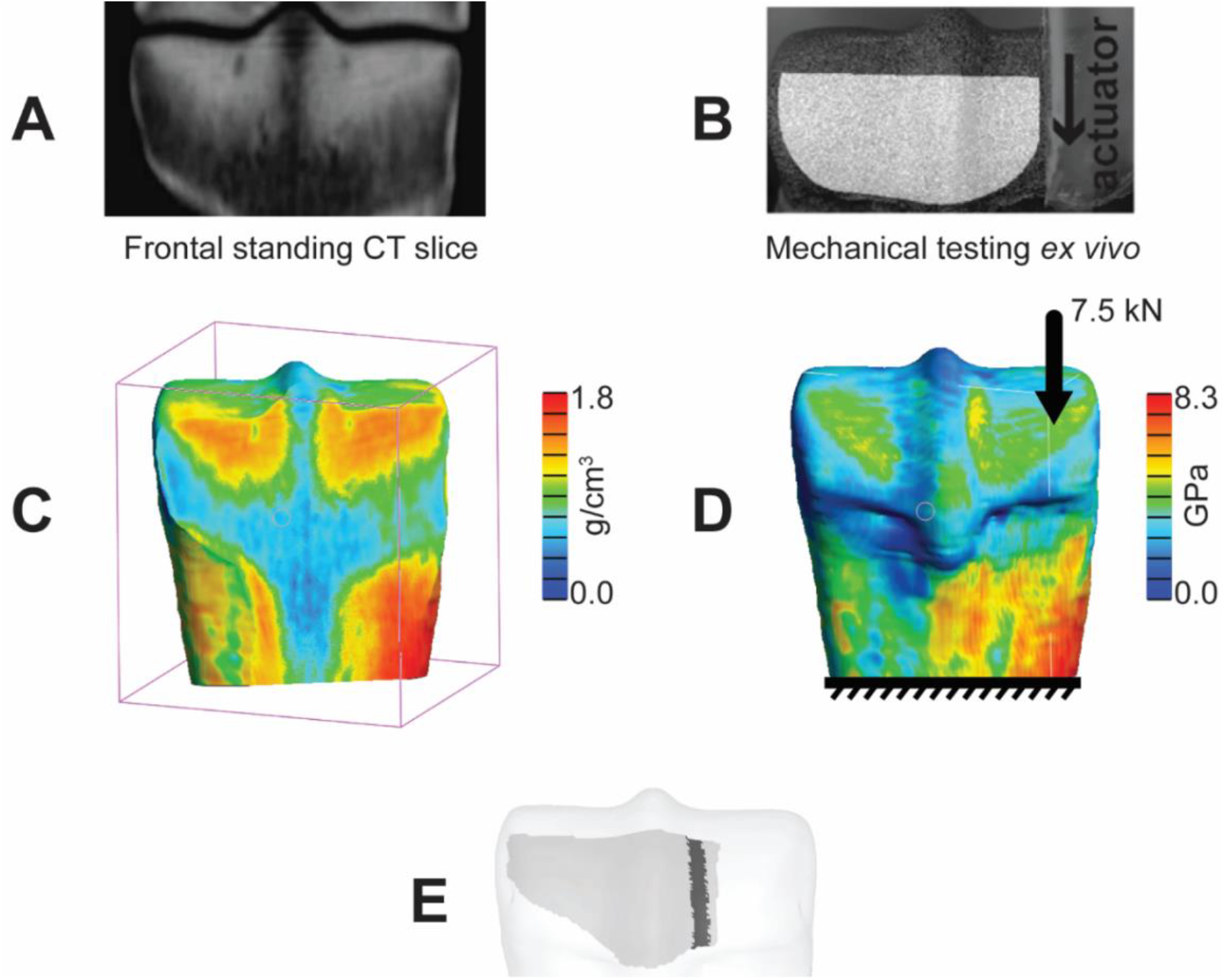
Finite element (FE) modeling pipeline. **A)** Standing CT (sCT) imaging was used for segmentation of the distal MC3 volume. **B)** Surface strain data from previous *ex vivo* testing data was used as the reference data for comparison against the FE results. **C)** Element-wise heterogenous HA density was captured from the sCT image set. **D)** Young’s modulus was assigned to individual elements and the *ex vivo* loading and boundary conditions were replicated in the model. **E)** The surface for which *ex vivo* mechanical testing data was available for (light gray) and the associated PSG region (dark grey) were registered on the 3D segmented bone end for comparison of the surface strain.

### 2.6. Model validation

The optimal *A_SCL_* and *D_LYS_* were used to model the other half of the CTRL and PSG-SBI condyles to validate the modeling approach. Predicted average PSG max principal strains were compared against the experimental results as the gold standard to determine the accuracy of the predicted PSG strain by the tuned virtual mechanical testing pipeline. Predictions of the tuned model were compared with the non-tuned model where *A*_*SCL*_ and *D*_*LYS*_ are zero.

## 3. Results

### 3.1. Tuned adaptive factor for the sclerotic regions

The group of four CTRL condyles that included low and high *V*_*SCL*_ (CTRL-2 and CTRL-3) and *ex vivo* PSG strain (CTRL-1 and CTRL-5) were identified (**Table S1**). The optimal *A*_*SCL*_ was a unique value for each of these condyles, ranging from -1.0 to 0.5. Models of condyles with smaller *V*_*SCL*_ required a negative *A*_*SCL*_ for their strain predictions to match the *ex vivo* results, and condyles with larger *V*_*SCL*_ required a positive *A*_*SCL*_ to capture the higher degree of microdamage in the subchondral sclerotic bone that results in a compromised Young’s modulus. Additionally, condyles with smaller *V*_*SCL*_ were less sensitive to change of *A*_*SCL*_ (**Table S2**). To find the optimal *A*_*SCL*_ function, it was plotted against *V*_*SCL*_, which showed a positive correlation (**Figure 3**), and a linear regression was made which resulted in the following relationship:

**Figure 3.**
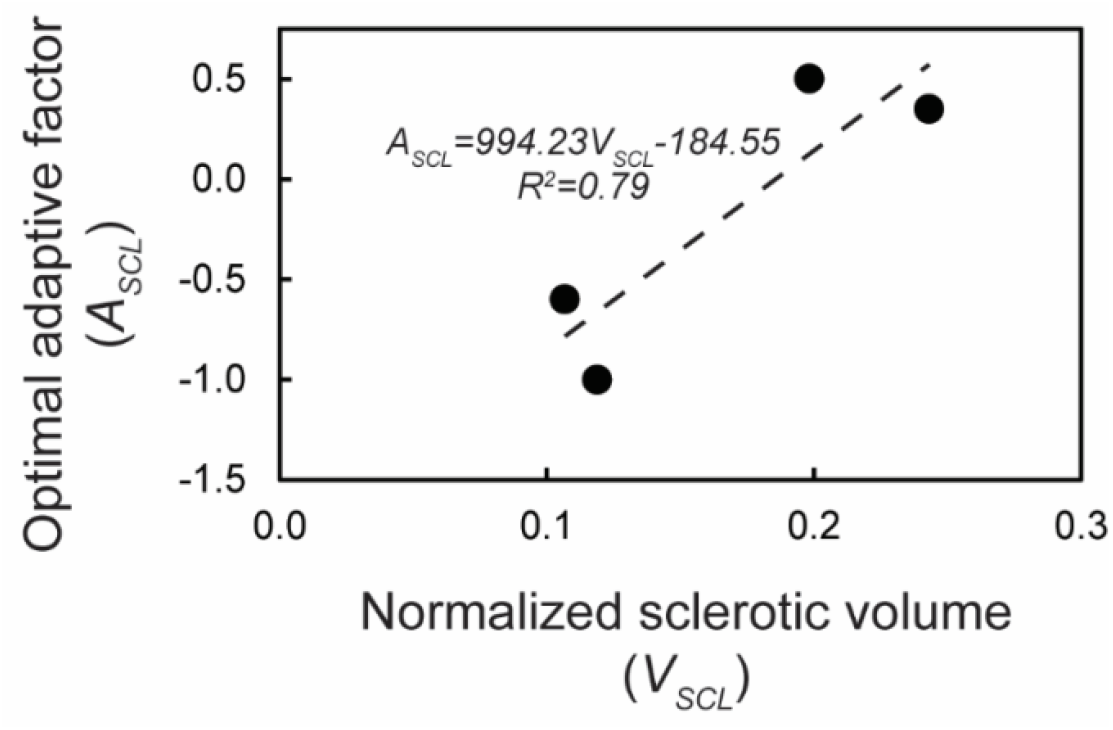
Adaptive factor *A*_*SCL*_ showed a positive relationship with the normalized subchondral sclerotic volume *V*_*SCL*_. This relationship was found by linear regression.

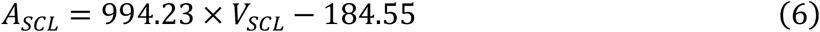

Equation (6) was used for the adoptive factor to adjust the Young’s modulus (Equation (4)) in the sclerotic regions in all the following models.

### 3.2. Tuned damage factor for the lytic regions

To select two of the PSG-SBI condyles for finding the optimal *D*_*LYS*_, PSG-SBI-3 was excluded due to its complex joint disease pathology that included palmar osteochondral disease (POD) as well as PSG-SBI. PSG-SBI-2 was excluded due to a very small overlap of the fatigue damage in the PSG area with the bone surface from which strain was measured [23]. Condyles PSG-SBI-1 and PSG-SBI-4 were selected and the optimal *D*_LYS_ for these condyles was found to be 0.80 and 0.50 respectively (**Table S3, S4**). As a result, an average constant value of 0.65 was determined as the overall optimal *D*_LYS_.

### 3.3. Model validation

The other half of the condyles were modeled to investigate the validity of the PSG strain predictions (**Table S5, Figure 4**). The optimal *A*_*SCL*_ and *D*_*LYS*_ determined in sections 3.1 and 3.2 were used to assign the Young’s modulus to the sclerotic regions and the lytic regions if present.

**Figure 4.**
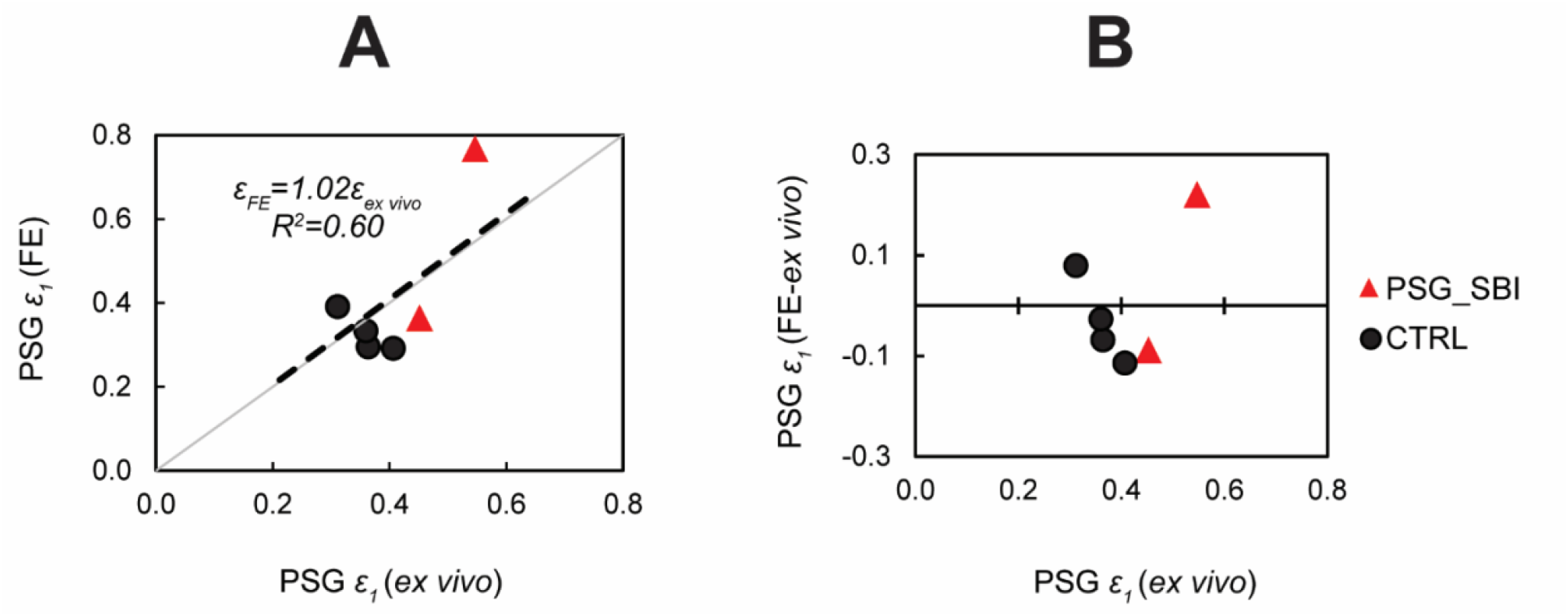
Comparison of the FE-predicted and the *ex vivo*-measured average PSG strain. Triangular and circular datapoints represent the PSG-SBI and CTRL condyles, respectively. **(A)** The FE pipeline predicts PSG strain with about 2% error on average. **(B)** Bland-Altman plot shows the slightly underestimated PSG strain by the FE pipeline except for the condyle PSG-SBI-3.

Tuning of Young’s modulus improved predictions of PSG strain over the non-tuned model (**Figure S3**). Predicted PSG strain matched *ex vivo* strain well (regression slope=1.02, R^2^=0.60, **Figure 4A**). The difference between FE-predicted and *ex vivo* PSG strain was smaller than 0.11 except for PSG-SBI-3 (**Table S5, Figure 4B**), which also had POD.

Surface strain fields were compared between *ex vivo* and FEA and showed good qualitative agreement (**Figure 5, Figure S4**). PSG-SBI-3 showed maximum principal strain values exceeding 1% in some regions of the PSG closer to the lysis. In this condyle, strain concentration was observed on the joint surface over the dorsal edge of the lytic volume. There was no sign of high strain concentration in the PSG of the CTRL condyles used for the model validation investigation. Overall, condyles with PSG-SBI showed higher PSG strain by qualitative comparison (**Figure S4**).

**Figure 5.**
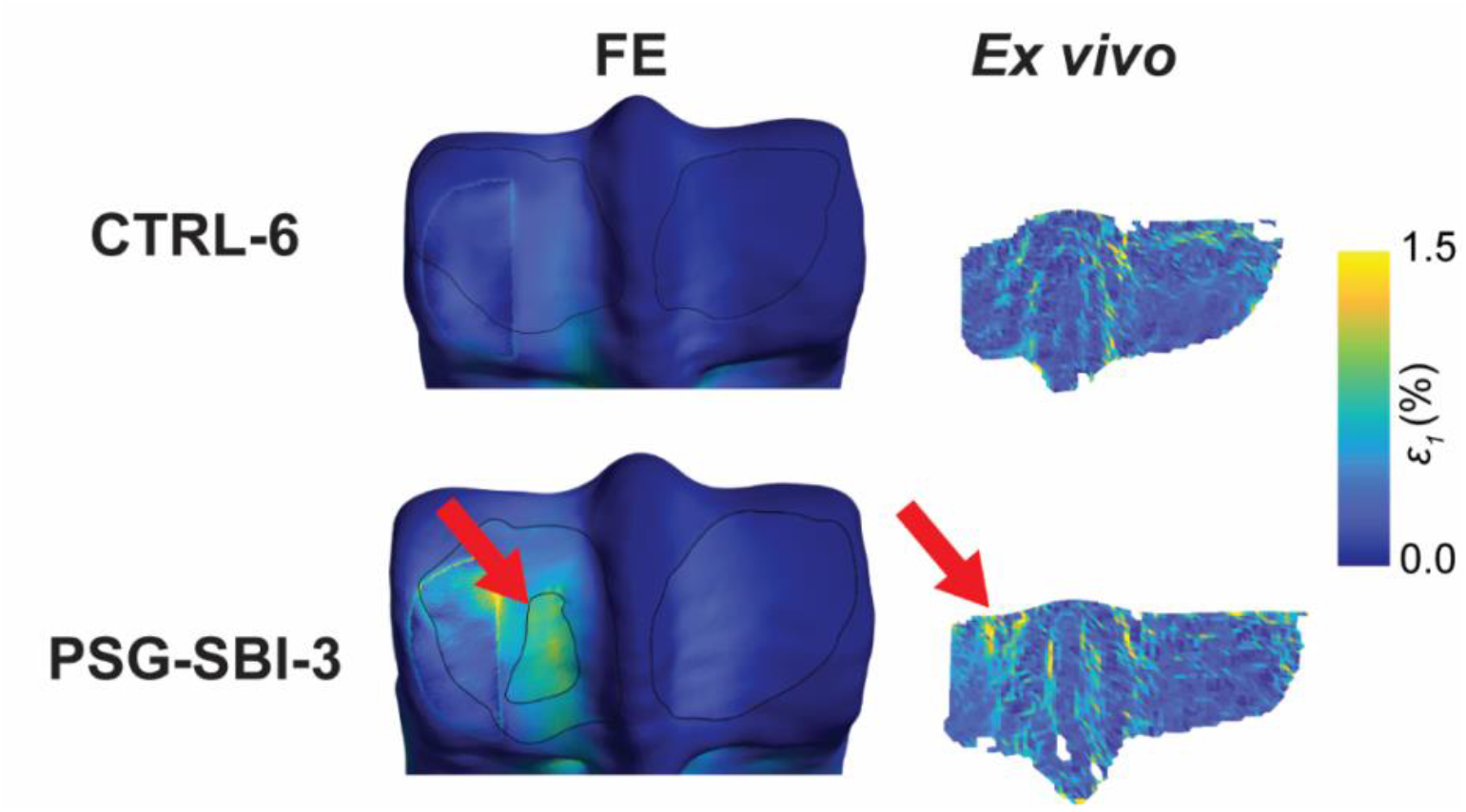
*Ex vivo*-measured and FE-predicted surface strain for the condyles CTRL-6 and PSG-SBI-3. *Ex vivo* strain data was not available for the dorsal aspect of the condyles as well as the area under the indenter. Red arrows show the location of the strain concentration in the PSG in the vicinity of the lysis that matches between the *ex vivo* and the FE-predicted results.

## 4. Discussion

The virtual mechanical testing pipeline presented here is the first model to our knowledge that can predict subject-specific load-induced strain on the joint surface of the distal MC3 bone in Thoroughbred racehorses using sCT-based FEA, validated against *ex vivo* mechanical testing data. While sCT imaging has revolutionized racehorse screening by making it practical for routine injury prevention screening, the quantified objective prediction of the biomechanical behavior of the subchondral bone provides valuable additional information that has the potential to assist clinicians in interpreting fatigue-induced subchondral structural changes in the fetlock and prevent potential underestimation of condylar stress fracture risk [25]. With growing use of sCT, longitudinal screening of horses is now feasible enabling assessments of lesion progression or healing of PSG SBI.

Tuning the Young’s modulus in the subchondral sclerotic and damaged bone resulted in three major improvements compared to the non-tuned model that assigns the same density-modulus relationship for the whole distal MC3. It improved prediction of PSG strain, explained a larger variation in the PSG strain measured *ex vivo*, and decreased the bias toward under-predicting PSG strain in bones with PSG-SBI. It is critical for this virtual mechanical testing pipeline to properly predict the high PSG strain in bones with PSG-SBI that were measured *ex vivo* [23], to prevent false negative diagnosis of Thoroughbreds that are at a potentially higher risk of condylar stress fracture when the goal of subject screening is to minimize risk of serious injury to both the Thoroughbred horses and their jockeys.

The subchondral sclerotic volume was used as a biomarker for the extent of fatigue damage in the sclerotic bone in our approach, since microdamage is not captured within the resolution of clinical CT. Model tuning indicated that the subchondral sclerotic bone volume is positively associated with fatigue-induced mechanical compromise in this region. This observation is aligned with previous findings that showed a positive association between bone volume fraction in the PSG and the degree of microdamage in this region [35].

Accurate prediction of bone strain is contingent on using a suitable density-modulus relationship, especially in the subchondral sclerotic bone. There are a limited number of studies that provide density-modulus relationship for the equine MC3 at the macro scale [34,36]. However, neither provide data on the subchondral sclerotic bone. Establishing a specific density-modulus relationship for this region has the potential to improve strain prediction accuracy. Despite the lack of such a site-specific relationship, the Young’s modulus modified by defining adaptive and damage factors in the present study could partially fill this gap in knowledge and successfully increased accuracy of bone strain prediction.

While the presented study provides a platform for biomechanical assessment within the quasi-static elastic deformation, future investigations on yield strength, post-yield behavior, and ultimate strength of MC3 equine bones, especially in the subchondral sclerotic region, could provide more insight into their plastic behavior and toughness [37]. The subchondral sclerotic MC3 bone, with bone volume fractions similar to cortical bone, loses its energy dissipation capabilities moving further from the joint surface despite having higher stiffness [38]. Although cortical bone has significantly lower toughness than trabecular bone [39], the difference in energy dissipation capabilities between sclerotic and non-sclerotic equine subchondral MC3 bone is a current gap in knowledge. Such information could improve prediction of stress fracture initiation and may enable prediction of its propagation path [40].

The main limitation of this study is the small number of samples with *ex vivo* mechanical testing data available for tuning and validation, mainly due to the burden of such experiments [23]. This forced tuning of the damage factor in the lytic regions to a constant value, while the association between the optimal damage and potentially some anatomical features such as the lytic volume could have been investigated, like what was performed for tuning the adaptive factor for the sclerotic bone. Another limitation is not including a bone with isolated palmar osteochondral disease (POD) in this study. Bones with POD have more extensive subchondral sclerosis [23,41–43] that results in larger normalized sclerotic volume. Condyle PSG-SBI-4 had both POD and PSG SBI pathologies with a high sclerotic volume. This resulted in the highest allowed adaptive factor (the highest allowed reduction in the Young’s modulus) for the sclerotic region, which could have caused the overestimation of PSG strain, and the highest error compared to *ex vivo* strain data. One limitation pertaining to the methodology presented here is the definition of homogenous adaptative and damage factors in the sclerotic and lytic regions, while there are higher degrees of microdamage closer to the articular surface, but such data are not available due to the resolution of clinical sCT. Future studies focused on spatial quantification of microdamage within these regions and its association with other fatigue biomarkers such as the normalized sclerotic volume can help alleviate this challenge.

## 5. Conclusion

In conclusion, the validated virtual mechanical testing pipeline presented here predicts subchondral MC3 PSG strain on a subject-specific basis and is a valuable objective mechanical assessment tool that is expected to improve identification of racing Thoroughbreds with high risk of condylar stress fracture. When applied to populations of Thoroughbred racehorse, this screening approach has the potential to reduce the incidence of serious or fatal injury in horses and produce an associated reduction in serious injuries to racehorse jockeys. This translational knowledge is also expected to advance stress fracture injury prevention in human athletes.

## Author Contributions

**Soroush Irandoust:** Conceptualization, Methodology, Software, Formal analysis, Investigation, Writing -Original Draft, Writing -Review & Editing, Visualization. **R. Christopher Whitton:** Conceptualization, Writing -Review & Editing, Funding acquisition. **Corinne R. Henak:** Conceptualization, Methodology, Formal analysis, Investigation, Resources, Writing -Original Draft, Writing -Review & Editing, Supervision, Funding acquisition. **Peter Muir:** Conceptualization, Methodology, Formal analysis, Investigation, Resources, Writing -Original Draft, Writing -Review & Editing, Supervision, Project administration, Funding acquisition.

## Acknowledgements

The authors would like to thank Faith Franseen, Christina Stevenson, and Nicola Brown for their help with image segmentation, and Dr. David Ergun for his help with acquiring the sCT image sets.

