## Supplementary Document for "Tuning and validation of a virtual mechanical testing pipeline for condylar stress fracture risk assessment in Thoroughbred racehorses"

Manuscript for the *Journal of the Royal Society Interface*

October, 2024

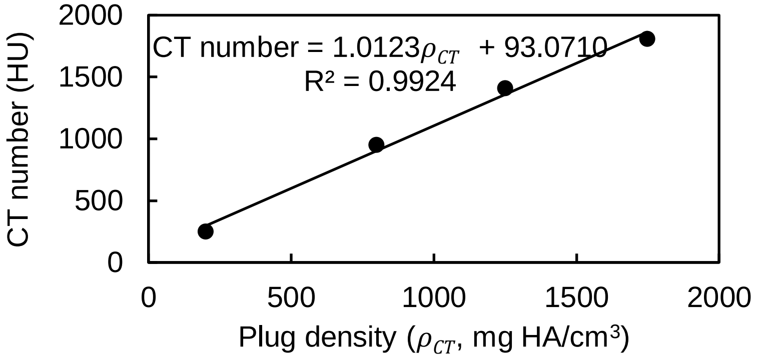


**Figure S1**. Calibration of the CT number by linear regression onto phantom plug density.

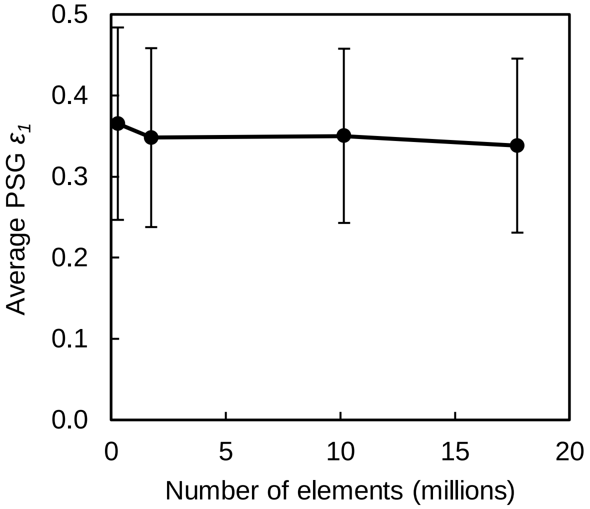


**Figure S2.** Mesh convergence analysis showing average PSG maximum principal strain for the condyle CTRL-5 for the four different mesh sizes modeled. Element edge lengths of 4.00-1.00 mm, 2.00-0.50 mm, 1.00-0.25 mm, and 0.80-0.20 mm were used in the sclerotic and distal MC3 regions, respectively. Error bars show standard deviation. Relative errors between consecutive mesh sizes were 4.9%, -0.6%, and 3.6% from left to right. The model with 17 million elements took 6.5 hours to run on a server with 192 processing cores and required around 400 GB of RAM memory. The selected mesh density, the model with 10 million elements took 4 hours to run on the same server and required around 150 GB of RAM memory. Higher number of elements could not be modeled due to increasing required memory. A model with 0.500-0.125 mm element edge lengths was created that resulted in more than 59 million elements and required more than 2 TB of RAM memory to run, but authors did not have access to a machine with such configuration.

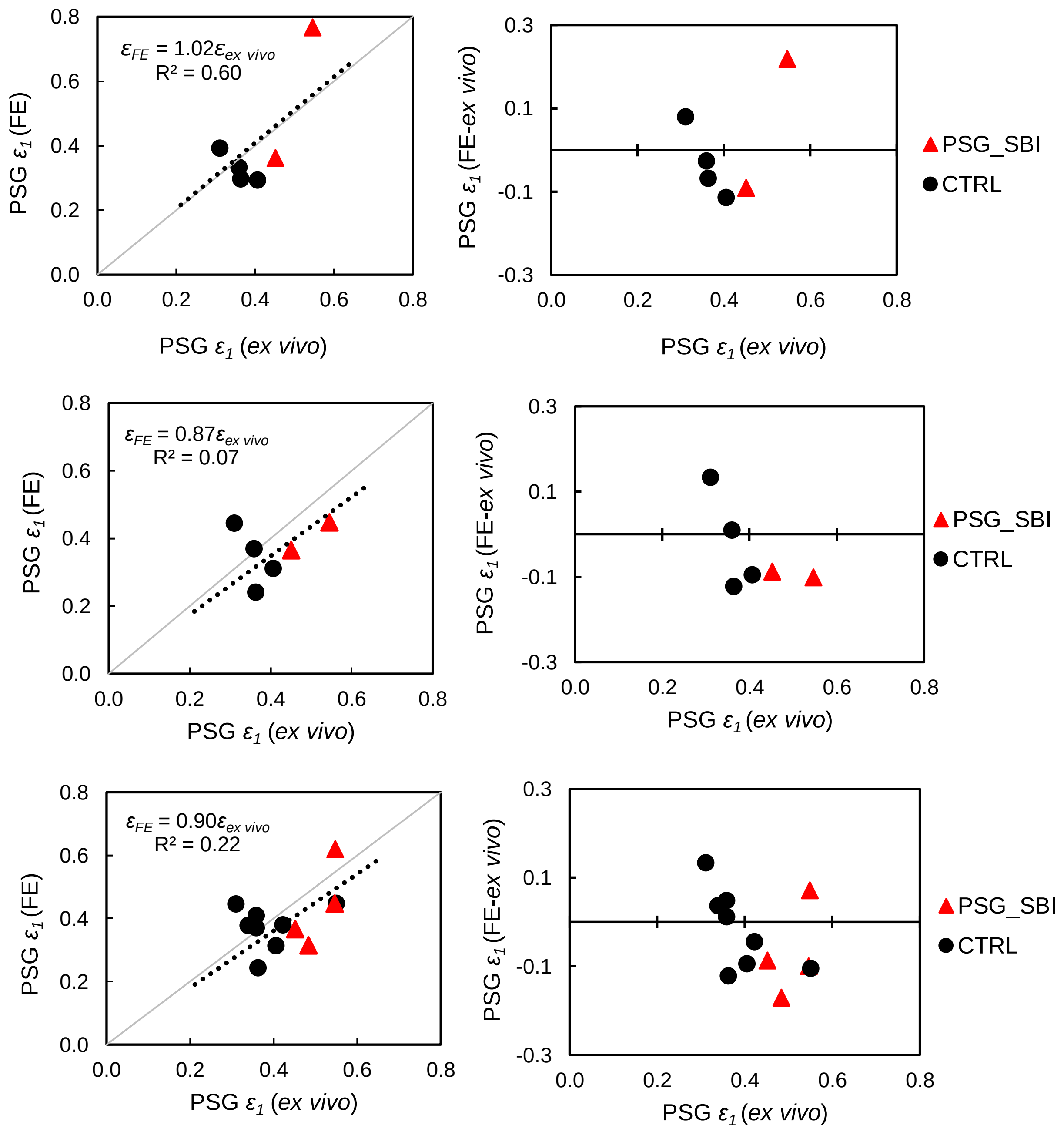


**Figure S3.** Comparison of the tuned model (first row) with non-tuned model (second and third rows). First and second rows show the data for only the specimens used for validation. The non-tuned pipeline underestimates PSG strain in bones with PSG SBI and for all specimens as well as the regression slope is less than 1, and does not explain the variance in the experimental data. All these limitations were resolved by the tuned modeling pipeline.

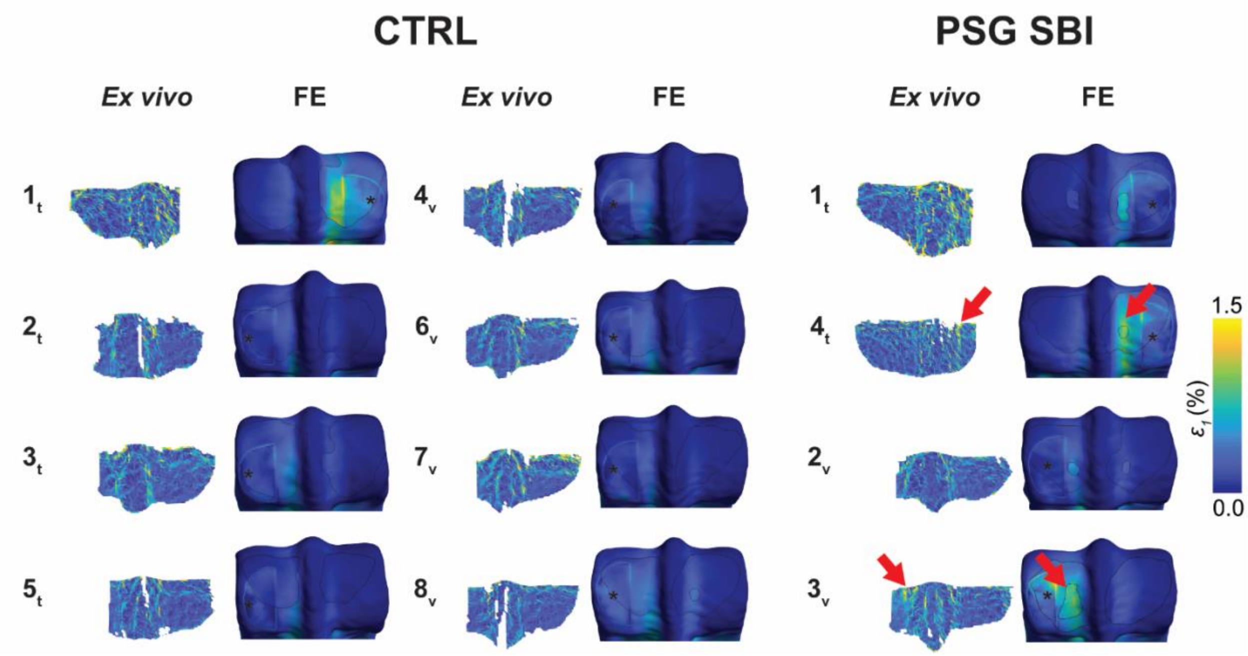


**Figure S4.** Maximum principal strain on the joint surface measured *ex vivo* and predicted with the developed virtual mechanical testing pipeline. Loaded condyles are marked with a star. Subscripts "t" and "v" indicate limbs used for training and validation respectively. Red arrows show elevated PSG strain colocalized with PSG fatigue injury. Higher PSG strain measured experimentally was predicted by the FE analysis as well.

**Table S1**. Normalized sclerotic volume $V_{SCL}$ (volume of the sclerotic region divided by the volume of the condyle), and the optimal $A_{SCL}$ in the tuning group.

| **Condyle** | $V_{SCL}$ | **Optimal** $A_{SCL}$ | **Average PSG** $\varepsilon_1$ (*FE*) | **Average PSG** $\varepsilon_1$ (*ex vivo*) |
|---|---|---|---|---|
| **CTRL-1** | 0.1984 | 0.50 | 0.5371 | 0.5513 |
| **CTRL-2** | 0.1068 | -0.60 | 0.3594 | 0.3599 |
| **CTRL-3** | 0.2431 | 0.35 | 0.4230 | 0.4236 |
| **CTRL-5** | 0.1188 | -1.00 | 0.3503 | 0.3403 |

**Table S2.** FE predicted average PSG maximum principal strain in the CTRL group for all values of $A_{SCL}$. Blank cells represent the $A_{SCL}$ values that were not modeled since they were determined not to be the optimal adaptive factor by inspection.

| $A_{SCL}$ | **CTRL-1** ($\varepsilon_{ex\ vivo}$ = **0.5513**) | **CTRL-2** ($\varepsilon_{ex\ vivo}$ = **0.3599**) | **CTRL-3** ($\varepsilon_{ex\ vivo}$ = **0.4236**) | **CTRL-5** ($\varepsilon_{ex\ vivo}$ = **0.3403**) |
|---|---|---|---|---|
| **0.50** | 0.5371 | 0.4887 | 0.4552 | 0.4107 |
| **0.45** | - | - | - | - |
| **0.40** | - | - | 0.4324 | - |
| **0.35** | 0.4993 | 0.4557 | 0.4230 | 0.3963 |
| **0.30** | 0.4893 | 0.4469 | 0.4146 | 0.3925 |
| **0.25** | 0.4803 | 0.4389 | 0.4070 | 0.3891 |
| **0.20** | 0.4721 | 0.4315 | 0.4002 | 0.3860 |
| **0.15** | 0.4645 | 0.4246 | 0.3939 | 0.3831 |
| **0.10** | 0.4576 | 0.4183 | 0.3881 | 0.3805 |
| **0.05** | 0.4511 | 0.4123 | 0.3828 | 0.3781 |
| **0.00** | 0.4452 | 0.4068 | 0.3779 | 0.3758 |
| **-0.05** | - | 0.4015 | - | 0.3737 |
| **-0.10** | - | 0.3966 | - | 0.3718 |
| **-0.15** | - | 0.3920 | - | 0.3700 |
| **-0.20** | - | 0.3876 | - | 0.3683 |
| **-0.25** | - | 0.3835 | - | 0.3667 |

| | | | | |
|---|---|---|---|---|
| **-0.30** | - | 0.3795 | - | 0.3651 |
| **-0.35** | - | 0.3758 | - | 0.3637 |
| **-0.40** | - | - | - | - |
| **-0.45** | - | - | - | - |
| **-0.50** | - | 0.3655 | - | 0.3599 |
| **-0.55** | - | 0.3624 | - | - |
| **-0.60** | - | 0.3594 | - | - |
| **-0.65** | - | 0.3566 | - | - |
| **-0.70** | - | 0.3538 | - | - |
| **-0.75** | - | 0.3512 | - | 0.3546 |
| **-0.80** | - | - | - | 0.3537 |
| **-0.85** | - | - | - | 0.3528 |
| **-0.90** | - | - | - | 0.3519 |
| **-0.95** | - | - | - | 0.3511 |
| **-1.00** | - | - | - | 0.3503 |

**Table S3**. Normalized sclerotic volume $V_{SCL}$ (volume of the sclerotic region divided by the volume of the condyle) for the medial and lateral condyles and the associated $A_{SCL}$ from equation (6), and the optimal $D_{LYS}$ in the tuning group.

| **condyle** | **$V_{SCL}$ (Medial, Lateral)** | **$A_{SCL}$ (Medial, Lateral)** | **Optimal $D_{LYS}$** | **Average PSG $\varepsilon_1$ (*FE*)** | **Average PSG $\varepsilon_1$ (*ex vivo*)** |
|---|---|---|---|---|---|
| **PSG-SBI-1** | 0.2291, 0.1780 | 0.4327, -0.0755 | 0.80 | 0.4755 | 0.4844 |
| **PSG-SBI-4** | 0.1513, 0.2378 | -0.3410, 0.5000 | 0.50 | 0.7881 | 0.5485 |

**Table S4.** FE predicted average PSG maximum principal strain in the PSG-SBI group for all values of $\boldsymbol{D_{LYS}}$. Blank cells represent the $D_{LYS}$ values that were not modeled since they were determined not to be the optimal damage factor by inspection.

| $\boldsymbol{D_{LYS}}$ | **PSG-SBI-1** ($\varepsilon_{ex\ vivo}$ = **0.4844)** | **PSG-SBI-4** ($\varepsilon_{ex\ vivo}$ = **0.5485)** |
|---|---|---|
| **0.85** | 0.4916 | - |
| **0.80** | 0.4775 | - |
| **0.75** | 0.4658 | 0.8375 |
| **0.50** | 0.4278 | 0.7881 |

**Table S5**. Normalized sclerotic volume $V_{SCL}$ for the medial and lateral condyles and the associated $A_{SCL}$ from equation (6), the optimal $D_{LYS}$ for the lytic volume if present, and the resulting predicted PSG strain compared to those measured *ex vivo* in the validation group.

| **Condyle** | $V_{SCL}$ **(Medial, Lateral)** | $A_{SCL}$ **(Medial, Lateral)** | $D_{LYS}$ | **Average PSG** $\varepsilon_1$ **(*FE*)** | **Average PSG** $\varepsilon_1$ **(*ex vivo*)** |
|---|---|---|---|---|---|
| **CTRL-4** | 0.2289, 0.1781 | 0.4303, -0.0745 | - | 0.2955 | 0.3637 |
| **CTRL-6** | 0.1797, 0.1087 | -0.0584, -0.7644 | - | 0.3907 | 0.3114 |
| **CTRL-7** | 0.1485, 0.1100 | -0.3693, -0.7516 | - | 0.3329 | 0.3597 |
| **CTRL-8** | 0.1513, 0.2378 | -0.3410, 0.5186 | - | 0.2920 | 0.4063 |
| **PSG-SBI-2** | 0.1713, 0.2163 | -0.1420, 0.3045 | 0.65 | 0.3614 | 0.4518 |
| **PSG-SBI-3** | 0.5816, 0.2422 | 0.5000, 0.5000 | 0.65 | 0.7655 | 0.5464 |
